# High-temporal resolution of microbial food web dynamics and structure during phytoplankton blooms in the Baltic Sea

**DOI:** 10.1101/2025.09.15.676335

**Authors:** Sohrab Khan, Klaudia Wdówka, Joanna Całkiewicz, Krzysztof Rychert, Lidia Nawrocka, Anetta Ameryk, Tanja Shabarova, Indranil Mukherjee, Karel Simek, Aneta Jakubowska, Mariusz Zalewski, Kasia Piwosz

## Abstract

Heterotrophic nanoflagellates (HNF) are a key component of the microbial food webs, playing an essential role in nutrient recycling and energy transfer in aquatic ecosystems. They have been typically considered to be bacterivores, but they can be also omnivorous (feeding on prokaryotes and other eukaryotes) and predatory grazers (feeding on other eukaryotes). Here, we combine CARD-FISH with both short and long-amplicon sequencing to resolve dynamics of key HNF groups during two high-frequency sampling campaigns in spring (March-May) and autumn (September-November) phytoplankton blooms in the coastal waters of the Baltic Sea. This approach allowed us to resolve the microbial food web dynamics within HNF communities at the phylotype level at time scales relevant to HNF duplication times. Omnivorous katablepharids and predatory MAST-2 dominated the HNF community, especially in spring. Bacterivorous groups (e.g., MAST-1, CRY1) were less abundant. Long-read sequencing revealed distinct seasonal shifts in dominant phylotypes, with Katable*pharis* sp. and MAST-2D peaking in spring, while other lineages became more prominent in summer and autumn. The high abundance of omnivorous HNF, compared to bacterivores, highlights their key role both as grazers of bacteria and flagellates and as a food source for predatory and omnivorous ciliates.

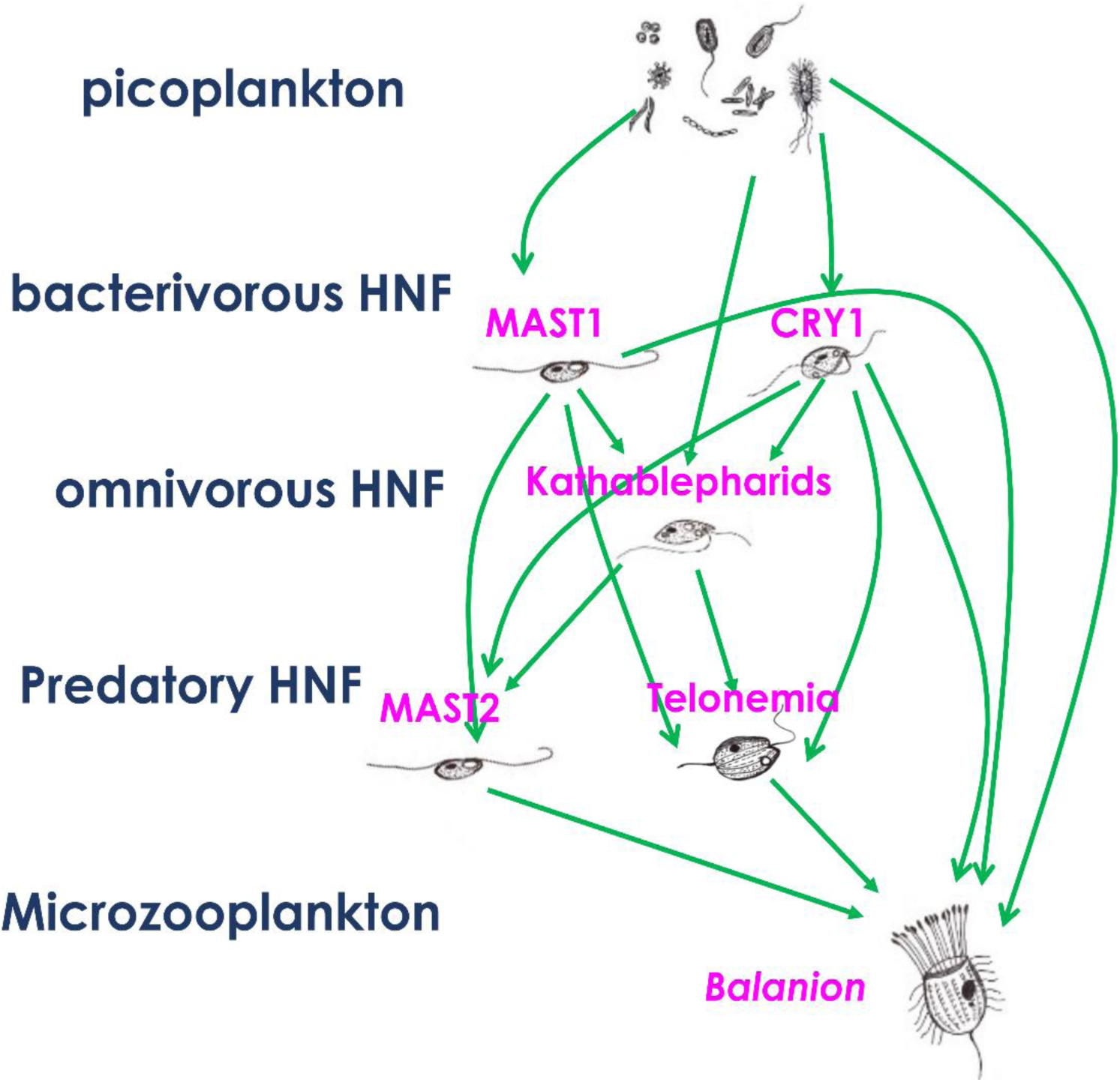

## Introduction

The phytoplankton blooms are key events in temperate regions, providing 90% of primary production for the marine food web at the global scale (Beltran-Perez and Waniek, 2022). Their seasonal dynamics and species succession are controlled by a combination of bottom-up environmental factors, such as light and nutrient availability, and top-down control factors, such as microzooplankton (*i.e.* large flagellates, ciliates and rotifers; Mironova et al., 2008), and mesozooplankton grazing (Mozetič et al., 2012). Additionally, temperature and salinity also influence these dynamics, playing important roles in shaping the brackish ecosystems. In the Baltic Sea, spring blooms are frequently dominated by diatoms, dinoflagellates and the prevalently autotrophic ciliate *Mesodinium rubrum* (Hjerne et al., 2019) that accounts for a substantial part of the annual primary production (Johansson et al., 2004). In autumn, diatoms, dinoflagellates, cryptophytes, and cyanobacteria dominate, reflecting seasonal shifts in phytoplankton composition (Wasmund and Uhlig, 2003). During the productive seasons, 16% to 50% of primary production is utilized by microbial food web (Witek et al., 1997). While seasonal fluctuations are well known to affect the Baltic’s microbial communities, the composition and dynamics of these communities, including heterotrophic protists which play key roles in nutrient cycling and energy transfer within the microbial food web, remain poorly understood. This is primarily due to challenges in visualizing and identifying small eukaryotic microbes. Understanding the community composition of heterotrophic protists is essential for gaining a comprehensive understanding of microbial interactions and their role in ecosystem functions. Heterotrophic nanoflagellates (HNF) are considered grazers of picoplankton (mainly bacteria) that form a link to higher trophic levels by feeding on it and being consumed by microzooplankton. Microzooplankton, in turn, are grazed by larger zooplankton, driving the carbon flow to higher trophic levels (Šimek et al., 1999; Jürgens and Jeppesen, 2000), thereby closing the microbial loop (Pernthaler and Amann, 2005; Kwon et al., 2017). By selectively grazing on picoplankton, HNF regulate picoplankton abundance and community composition (Simek et al., 1997; Jezbera et al., 2006; Jeuck and Arndt, 2013; Grujčić et al., 2015).

Recently, an extended model of the aquatic microbial food web described new, non-linear trophic interactions within nanoplankton, providing a novel perspective on the roles of HNF (Piwosz et al., 2021). Prokaryotes are grazed by small bacterivorous flagellates, but they can also be consumed by omnivorous flagellates and small ciliates. Larger omnivorous/predatory flagellates, which feed on both bacteria and bacterivorous flagellates, are grazed by predatory ciliates (Šimek et al., 2020). Meanwhile, small ciliates, aside from being predators, also consume bacteria. These flagellates and ciliates are preyed upon by microzooplankton, which can also feed on smaller bacterivorous flagellates, making the microbial food webs considerably more complex than assumed in the classical description of the microbial loop concept (Azam et al., 1983). The ecological importance of omnivorous and predatory HNF (Šimek et al., 2020; Piwosz et al., 2021) remains far less explored compared to bacterivorous flagellates.

The size distribution of the HNF community affects the food web structure, with smaller HNF (< 3 µm) being bacterivorous, while larger ones (3-8 µm) exhibiting omnivorous or predatory behaviour (Arndt et al., 2000; Jürgens and Massana, 2008). The small size and lack of distinctive morphological features make HNF challenging to identify using standard microscopic analysis. Additionally, HNF communities in the Baltic Sea are influenced by environmental gradients at both large and local scales (Hu et al., 2016; Piwosz et al., 2018). Recent findings, including our data (Piwosz et al., 2025), shows that 80% of HNF are smaller than 5 µm, with picoplanktonic HNF (< 3 µm) making up over 37% of the population, and only 2% exceeding 10 µm. This indicates that most HNF may be omnivorous and/or predatory, as proposed in the new model (Piwosz *et al*., 2021).

The aim of the study was to provide deeper insight into the dynamics of grazers’ levels within the microbial food webs and to understand how seasonal changes affect microbial community composition. While high-throughput sequencing of environmental DNA has greatly advanced our understanding of the diversity of microorganisms in various environments (Del Campo et al., 2018), especially community-level processes (Logares et al., 2013; Villarino et al., 2018; Santoferrara et al., 2020), it does not provide insights into the abundance, morphology, trophic roles or feeding preferences of the identified groups (Piwosz et al., 2020; Mukherjee et al., 2024). To overcome these identification and quantification challenges, we utilized group-specific CARD-FISH probes and sequencing techniques. CARD-FISH enables precise visualization of target HNF groups, addressing size and morphological limitations, while sequencing provides a detailed view of community composition and diversity. We conducted a high frequency sampling (three-times a week) to match protistan growth rates yielding also short-lived peaks of small protists, since the typical doubling time of prokaryotes and protists in the epilimnetic waters ranges from 12 hours to days (Zeder et al., 2009; Eckert et al., 2012; Šimek et al., 2018). This combination of approaches allows for a full understanding of HNF dynamics and their trophic roles in the microbial food webs. We focused on several groups of heterotrophic nanoflagellates that play distinct roles in the microbial food web, including ubiquitous bacterivores (MAST-1, MAST-4, CRY1 lineage of cryptophytes and *Paraphysomonas* spp.) (Massana et al., 2006b; Piwosz et al., 2016; Šimek et al., 2022; Šimek et al., 2023), omnivorous katablepharids, and predatory MAST-2 and Telonemia (Massana et al., 2006b; Grujcic et al., 2018; Boukheloua et al., 2024). Our study reveals seasonal shifts in HNF communities in the Baltic Sea, driven by temperature and nutrients, and highlights their role in the microbial food webs.

## Experimental Procedures

### Sampling

Water samples were collected three time per week from a 8-meter-deep coastal station (54° 32’ 54.6972’’ N; 18° 34’ 2.6076’’ E) in the Gulf of Gdańsk (southern Baltic Sea, Poland) during the spring (27^th^ March to 4^th^ May) and autumn (18^th^ September to 20^th^ October) seasons of 2023. Additionally, monthly samples were collected on June 5th, July 4^th^, July 31^st^, and November 2^nd^, 2023. Ten litres of water were collected in a clean, detergent-washed container, rinsed three times with MilliQ and then with the sampled seawater. The samples were transported to the laboratory within one hour for further analysis. Depth profiles of water temperature and salinity were acquired *in situ* using a Sontek Castaway CTD probe (USA). Weather conditions (cloudiness, wind, waves, and air temperature) were also recorded. The collected seawater was divided into two fractions: unfiltered and 20-µm filtered. The filtered fraction passed through a 20-µm plankton net was used to analyse prokaryote and heterotrophic nanoflagellates abundance, and to acquire biomass for amplicon sequencing. The unfiltered fraction was utilized for phytoplankton, heterotrophic dinoflagellates and ciliate analyses, as well as for nutrient and chlorophyll measurements. A table containing all measurements is available as xlsx file at https://zenodo.org/ (DOI: 10.5281/zenodo.15411382).

### Abundance of Prokaryotes and Heterotrophic and Phototrophic Nanoflagellates

Three ml of filtered sample were fixed with sterile-filtered formalin (final concentration 2%) and filtered onto black-stained polycarbonate (PC) filters (25 mm diameter, 0.2 µm pore size). Once air-dried, the filters were stained with 4’,6-diamidino-2-phenylindole (DAPI; 1 μg ml^−1^) (Coleman, 1980). Cells were counted across thirty fields of view (FOVs) using fluorescent microscopy with UV/Blue light excitation/emission (Olympus BX50, 1000× magnification). To quantify heterotrophic and phototrophic nanoflagellates, 25–50 ml of water samples were fixed with alkaline Lugol’s solution (final concentration 1%) followed by the addition of formalin to a final concentration of 2%. Afterwards, the samples were decolorized using 3% Na_2_S_2_O_3_ (Sherr et al., 1987), filtered onto black-stained polycarbonate (PC) filters (pore size 0.8 μm, diameter 25 mm, Isopore, Millipore), air-dried, and stained with DAPI. At least 30 microscopic fields of view (FOVs) per sample were counted using fluorescent microscopy (Olympus BX50, 1000× magnification), with UV/blue light excitation/emission for DAPI signals and blue/red light excitation/emission for autofluorescence of chloroplasts of autotrophic cells. A table containing cell abundances is available as xlsx file at https://zenodo.org/ (DOI: 10.5281/zenodo.15411382).

### Microscopic Microozoo- and Microphytoplankton Analysis

Two water samples of 50 ml each were fixed with acidic Lugol’s solution (final concentration 1%). Both groups were analysed using method recommended by the Helsinki Commission for studies on phytoplankton (HELCOM 2010). Shortly, samples were concentrated in the Utermöhl chamber and observed under an inverted microscope. The list of currently accepted taxon names was compiled based on the World Register of Marine Species (Ahyong et al., 2023).

Ciliates were counted and measured under an inverted microscope (Nikon Eclipse TS100) at magnifications of 400×. Ciliates were identified to the lowest possible taxonomic level. Fixation with Lugol’s solution does not cause loss of ciliate cells but hinder their identification (Rychert et al., 2016). Phytoplankton was analysed under an inverted microscope (Delta Optical IB-100, Poland) at magnifications of 200× and 400×. The counting units (n) were cells, colonies, coenobia, or 100 µm long trichomes. To calculate the biomass (biovolume), the species were approximated to simple geometric or combined forms. Counting were performed following the recommendations of the Baltic Monitoring Programme (Napiórkowska-Krzebietke and Kobos, 2016; Jetoo and Tynkkynen, 2021).

### Chlorophyll *a* analysis

For chlorophyll *a* (Chl-*a*) measurements, 100 mL of the water sample was filtered onto GF/F filters (Whatman, UK), placed in the extraction tubes and immediately frozen at -20°C, before being analysed within a month. The filters were then extracted with 96% ethanol at room temperature for 12 hours. Chl-a concentrations were measured using a Turner Design fluorimeter (Fluorimeter 10-005, Canada) following the protocol recommended by the Helsinki Commission for the Baltic Sea (HELCOM 2010). The excitation wavelength was 425-430 nm, and the fluorescence emission wavelength was 663-672 nm. This method is based on the previous research (Holm-Hansen et al., 1965; Lorenzen, 1967; Jespersen, 1987). A table containing chlorophyll data is available as xlsx file at https://zenodo.org/ (DOI: 10.5281/zenodo.15411382).

### Nutrients

Water samples of 500 mL were collected in acid-cleaned bottles, frozen at -20°C, and analysed within a month. Concentrations of total nitrogen (N-tot), nitrate nitrogen (N-NO_3_), nitrite nitrogen (N-NO_2_), and ammonium nitrogen (N-NH_4_), collectively referred to as dissolved inorganic nitrogen (DIN), as well as phosphate phosphorus (P-PO_4_) and total phosphorus (P-tot), were determined following the method of Grasshoff et al.(Grasshoff et al., 2009). A table containing nutrients data is available as xlsx file at https://zenodo.org/ (DOI: 10.5281/zenodo.15411382).

### Catalysed Reporter Deposition – Fluorescence *in situ* Hybridization (CARD-FISH)

Based on the sequencing data (see below), we selected the following HNF lineages to determine their absolute abundances in the samples by CARD-FISH: MAST-2 (Massana et al., 2006b), MAST-4 (Massana et al., 2002), Katablepharidacea (Piwosz et al., 2025), MAST-1 (Massana et al., 2004), genus *Paraphysomonas* (Caron et al., 1999), phylum Telonemia (Mukherjee et al., 2024) and Cry1 lineage of cryptophytes (Piwosz et al., 2016). The CARD-FISH was done following the protocol described in Piwosz et al. (Piwosz et al., 2021). The hybridization conditions, and sequences of the probes utilized in this study are detailed in Supplementary Table 1. A 200-ml sample was fixed as described for the total HNF abundance. Fixed samples were stored in the dark at room temperature for 1 hour, filtered onto white polycarbonate filters (0.8-µm pore size, 47-mm diameter; Isopore, Millipore), rinsed with 30 ml of sterile MilliQ water, air dried, and then stored at –20°C until further processing.

A minimum of 30 microscopic fields of view (FOVs) per sample were counted to ensure a total of 100 DAPI-stained cells using fluorescent microscopy (Olympus BX50, 1000x magnification). The ratios of hybridized cells were counted in proportion to the number of HNF detected in DAPI. A table containing abundances of all HNF groups analysed by CARD-FISH is available as xlsx file at https://zenodo.org/ (DOI: 10.5281/zenodo.15411382).

### Cell size measurements

HNF cell sizes were measured following CARD-FISH hybridization with a general EUK-516 probe (Amann et al., 1990). Epifluorescence microscopy (AxioImager.M1, Carl Zeiss, Germany) was used to photograph at least 20 FOVs at 1000× magnification. Identification of HNF was based on the probe signal, DAPI signal, and the lack of chloroplast autofluorescence. A total of 1,700 cells were measured across all samples, with 50 cells analysed per sample using the length tool in Zen Pro Microscopy Software. A table containing cell-size data is available as xlsx file at https://zenodo.org/ (DOI: 10.5281/zenodo.15411382).

### DNA sample collection and extraction

Biomass for amplicon sequencing was collected by filtering 0.3 to 2.5 litres of 20-µm filtered water on nitrocellulose membrane filters (3-µm pore size, 47-mm diameter) under aseptic conditions. This approach was aimed to enrich the contribution of larger, presumably omnivorous and predatory HNFs (Piwosz et al. 2021) by reducing contribution from both picoeukaryotes, mostly bacterivorous HNF and green algae (Jürgens and Massana, 2008; Piwosz, 2019) and microeukaryotes, such as diatoms, dinoflagellates, ciliates and larger zooplankton. The filters were immediately frozen and stored at –80°C until further processing within a month. DNA extraction was performed using a custom, optimized protocol that combines the GeneMATRIX Soil DNA Purification Kit (Eurx, Gdańsk, Poland) and GeneMAGNET PCR Clean-Up Beads (Eurx). Extracted DNA samples were stored at -80°C until further analysis. The quality of the extracted DNA was assessed by gel electrophoresis, and the concentrations were measured with QubitFlex Fluorometer using Qubit 1x dsDNA HS Assay Kit (Invitrogen, Eugene, USA). To investigate the HNF community composition with both taxonomic resolution and broader diversity coverage, we applied a dual sequencing approach. Short amplicons (V6– V8 region of the 18S rRNA gene, primers 1183F and 1625R; (Latz et al., 2022)) were sequenced by Novogene (Cambridge, UK) on the Illumina NovaSeq platform (2 × 250 bp), offering high-throughput and fine-scale taxonomic profiling. To better resolve taxonomic composition of most abundant HNF groups, long amplicons (from V4 of the 18S rRNA to D9 of the 28S rRNA gene; primers V4F (Stoeck et al., 2010) and 21R (Schwelm et al., 2016)) were sequenced at Rush University Genomics and Microbiome Core Facility (Chicago, USA) using the PacBio Sequel II platform. Libraries were prepared using Fluidigm barcodes for equimolar sample representation and processed with the SMRTbell system, followed by sequencing on an 8M SMRT cell with a 30-hour movie runtime. This complementary strategy allowed us to obtain a more comprehensive view of HNF diversity and community structure.

Preventive measures were implemented to avoid contamination throughout the DNA extraction process. The extraction was conducted under a fume hood sterilized with a UV lamp to minimize cross-contamination, especially from airborne particles. All working surfaces and necessary equipment were regularly cleaned using boiling MilliQ water, followed by 70% alcohol and alkaline bleach to ensure a sterile environment. DNA from blank controls, prepared by filtering sterile MilliQ water, was extracted along with the samples. No DNA was detected in these blank controls. They were nevertheless sent for library preparation and sequencing, however, their amplification failed, confirming the absence of any contamination during biomass collection and DNA extraction.

### Short amplicons sequence analysis

The Illumina sequencing of the V6-V8 fragment of the 18S rRNA gene yielded a total of 2,145,661 reads across all 18S samples. Quality filtering of the raw sequence data was performed using the DADA2 pipeline in R (R 4.3.2) (Callahan et al., 2016). This approach removed low-quality reads, denoised the data, detected chimeras, and merged paired end reads, resulting in high-quality amplicon sequence variants (ASVs). Each sample had a minimum of 32,394 reads, with an average of 46,906 reads per sample. Chloroplast and non-eukaryotic sequences were filtered out to ensure precise analysis. Taxonomic classification of the sequence variants was performed using the DADA2 ‘assignTaxonomy’ function, with the PR2 database v5.0.0 (Guillou et al., 2012) as reference. Groups known to include HNF (phyla Katablepharidacea, Telonemia, Ancyromonadida, Apusomonada, Metamonada, Nibbleridia, Picozoa, Rhizaria, Rigifilida, Tubulinea, Opisthokonta; division Bigyra; families MAST-1, MAST-2,MAST-4, Basal_Cryptophyceae-1; order Paraphysomonadales) were further analysed for their relative abundances in the samples, in order to select target groups for CARD-FISH. Phyloseq objects containing data for all eukaryotic reads and reads coming from HNF groups are available as rds file at https://zenodo.org/ (DOI: 10.5281/zenodo.15411382).

### Long amplicons sequence analysis

Post-sequencing, data processing took place at the Rush University Research Bioinformatics Core Facility. Reads were demultiplexed and dereplicated using a DADA2-based pipeline optimized for long-amplicon sequencing (Callahan et al., 2019). Chimeric sequences were identified and removed using the BimeraDenovo algorithm, applying a minimum parent-over-abundance threshold of 3.5. Each sample had a minimum of 5,974 reads, with an average of 18,333 reads per sample. Taxonomic assignment was performed on 18S rDNA fragments extracted from nonchimeric ASVs with Barrnap software (https://github.com/tseemann/barrnap) using the PR2 database v5.0.0 (Guillou et al., 2012) as reference. A table containing sequence information, abundance and taxonomic assignment of all ASVs is available as xlsx file at https://zenodo.org/ (DOI: 10.5281/zenodo.15411382).

### Phylogenetic analysis

Phylogenetic tree analysis for katablepharids and MAST-2 was performed following the procedure optimised for the EUKARYOME database (Tedersoo et al., 2024). In addition to ASVs from long amplicon sequencing, the 18S sequences affiliated with both groups were retrieved from ENA database. All sequences were initially aligned using the MAFFT online tool with default settings (Katoh et al., 2019). The alignment was manually improved using ALIview (Larsson, 2014), and subsequently trimmed using ClipKIT (V.2.1.3) with the gappy parameter set to 0.9 (Steenwyk et al., 2020), resulting in the removal of 1.31% (57/4358) sites from the original MAST alignment and 5.59% (257/4600) sites from the katablepharids alignment. The phylogenetic trees by maximum likelihood was inferred using IQ-TREE with default setting and the TN+F+I+G4 model, automatically selected based on the Bayesian Information Criterion (Minh et al., 2020). The consensus tree was visualized using figTree (v.1.4.3.) and edited for final presentation with Inkscape (V.1.3.2). Alignments in fasta format and trees in Newick format are available at https://zenodo.org/ (DOI: 10.5281/zenodo.15411382).

### Statistical Analysis

The statistical relationships between total HNF abundance and abundances of the most abundant groups revealed by CARD-FISH analysis (katablepharids and MAST-2, hereafter referred to as biological variables) and the measured environmental factors (temperature, salinity, N-NH_4_, N-NO_3_, DIN, N-tot, P-PO_4_, P-tot and N-tot to P-tot (N:P) ratio, hereafter referred to as environmental variables) were calculated to assess seasonal patterns separately in spring (March 27 to May 4, 2023, 16 samples) and autumn (September 18 to October 20, 2023, 15 samples). All analyses were conducted in R using the dplyr (ver. 1.1.4), tidyr (ver. 1.1.3), ggplot2 (ver. 3.5.1), broom (ver. 1.0.6), and ggcorrplot (ver. 0.1.4.1), stats (ver. 4.4.0) packages. Calculations and visualizations were performed separately for spring and autumn. Pearson correlation coefficients (r) were calculated using the cor function, and significance was tested with cor.test (Student’s t-test). When multiple comparisons were performed, p-values were adjusted using the Holm–Bonferroni correction. To identify potential time-lagged dependencies between biological and environmental variables, cross-correlation function (CCF) analysis was conducted separately for each season. The maximum lag was set to one and two sampling intervals, with correlation significance assessed using p-values. The ccf function from the stats package in R was used, and results were visualized as CCF function plots. For each variable pair, CCF plots were generated, and correlation results were recorded for different lags. In selected cases, both zero-lag and time-lagged regressions were performed for visual inspection of relationship strength across different lags. The results were visualized as heatmaps using the ggcorrplot package. To minimize the confounding effect of temperature and to better uncover other underlying relationships, residual regression was applied. For each biological variable, residuals were extracted from linear models with temperature as the predictor (lm(value ∼ Temperature)), performed separately for spring and autumn. These residuals were then used in subsequent correlation analyses, effectively removing the direct influence of temperature, which is a typical driver of seasonal variation.

### Data availability

The sequences from Illumina and PacBio seqencing were deposited in the EMBL database under the BioProject PRJEB89350. Other data presented in this work are available at https://zenodo.org/ (DOI: 10.5281/zenodo.15411382).

## Results

### Environmental conditions

Water temperature varied seasonally, beginning at 4°C in early spring (March), and rising to 8.3°C by May. During summer (June to August), the average temperature reached 16.7°C. In autumn (September to November), temperature gradually declined from 19.5°C to 11.2°C (Supplementary Figure 1). Salinity remained mostly steady throughout the sampling period, varying between 6.0 and 7.5. DIN concentration varied seasonally, ranging from 1.04 to 7.29 µmol dm⁻³ in spring, 1.69 to 2.24 µmol dm⁻³ in summer, and 1.17 to 5.02 µmol dm⁻³ in autumn. Concentrations of N-tot showed two maxima in April, and they ranged from 6.85 to 30.20 µmol dm⁻³ in spring, were around 6 µmol dm⁻³ in summer, and varied from 3.21 to 9.88 µmol dm⁻³ in autumn. Concentrations of P-PO₄ increased from 0.18–1.12 µmol dm⁻³ in spring to 0.33–0.43 µmol dm⁻³ in summer and 0.43–0.82 µmol dm⁻³ in autumn. P-tot was highest in spring (1.08–13.33 µmol dm⁻³) and declined through summer (0.97–1.08 µmol dm⁻³) and autumn (0.92-1.77 µmol dm⁻³). N: P ratio was generally below 16 over the seasons (1.3-14.3) indicating nitrogen limitation, except for three days between 17 and 24 of April, when values from 18.7-23.4 suggested phosphorus limitation.

### Phytoplankton and Chlorophyll-*a*

Phytoplankton abundance and Chl-*a* concentrations exhibited closely linked seasonal patterns. Phytoplankton abundance ranged from 25 to 1050 cells ml^-1^, with the maximum recorded during the spring bloom in late April (Figure 1A). Concordantly, we observed high Chl-*a* concentrations from mid to the end of April, with the highest concentration exceeding 10 µg L⁻¹ (Figure 1B). The spring bloom was dominated by diatoms *Chaetoceros wighamii* and *Ch. danicus*, along with dinoflagellates such as *Dinophysis acuminata and* chrysophytes such as *Chrysidiastrum catenatum*. As summer season progressed, we observed a decrease in phytoplankton abundance ranging from 90 to 260 cells ml^-1^, while chlorophyll-a concentrations ranged from 3.6 to 5.4 µg L^-1^. Filamentous and potentially toxic cyanobacterial species *Aphanizomenon flos-aquae* and *Nodularia spumigena* dominated during summer, reflecting the intense cyanoblooms typical of the Baltic Sea, with Bacillariophyceae (*Cyclotella choctawhatcheeana*) and Dinophyta (*Dinophysis acuminata* and *D. norvegica*) being also present. In autumn, phytoplankton abundance varied between 1.4 and 380 cells ml^-1^, reaching its highest level in early October. Meanwhile, Chl-*a* concentrations peaked at 6.6 µg L⁻¹ in September before gradually declining to 1.3 µg L⁻¹ by the end of autumn. At that time, Bacillariophyceae (*Dactyliosolen fragilissimus* and *Centrales*) and Dinophyta remained abundant, with minor contribution of cryptophytes and chrysophytes.

**Figure 1.**
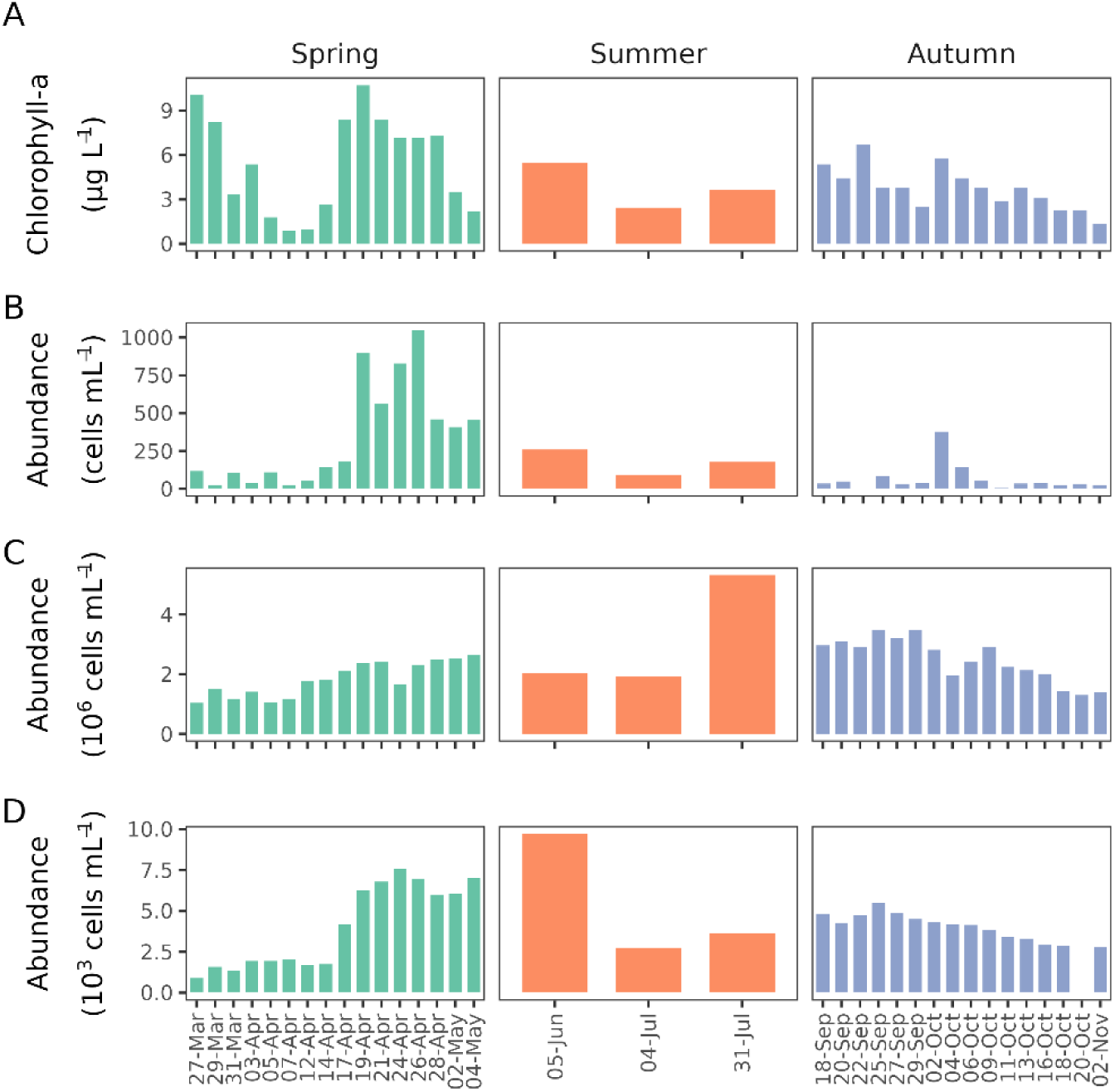
Chlorophyll-a concentration (A); phytoplankton abundance (B); bacteria abundance (C); and HNF abundance (D) during high-frequency sampling campaigns in spring and autumn, and in monthly summer samplings. Y-axis differs between the panels.

### Abundance of Prokaryotes and Heterotrophic Nanoflagellates

Prokaryote abundance ranged from 1.04 × 10^6^ to 5.31 × 10^6^ cells ml^-1^, peaking in July during high summer temperatures and was the lowest in March (Figure 1C). During spring (Mar-May), prokaryote numbers increased alongside rising temperatures, ranging from 1.04 × 10^6^ to 2.54 × 10^6^ cells ml^-1^. In autumn (Sep-Nov), as temperatures dropped, prokaryote numbers declined from 3.5 to 1.9 × 10^6^ cells ml^-1^.

HNF abundance ranged remarkably during the high-frequency sampling period. HNF numbers were the lowest in March with, around 900 cells ml^-1^ on the first sampling day and steadily rose reaching 7 × 10^3^ cells ml^-1^ in May as temperatures increased (Figure 1D). During summer, HNF abundance ranged from 2.76 × 10³ to 9.70 × 10³ cells ml⁻¹, with a peak observed in June. In autumn, HNF abundance gradually decreased from 5.50 × 10^3^ cells ml^-1^ in September to 2.77 × 10^3^ cells ml^-1^ in November.

### HNF community size structure

The size distribution of HNF showed conspicuous seasonal patterns (Figure 2). In spring, 67% of HNF were <5 µm, of which 32% were <3 µm, and only a small fraction (2.2%) exceeded 10 µm. During summer, the dominance of <5 µm HNF increased to 77%, of which 21% were <3 µm and less than 1% larger than 10 µm. In autumn, HNF remained predominantly <5 µm (77%), with 10.7% <3 µm and no significant presence of HNF larger than 10 µm (Figure 2).

**Figure 2.**
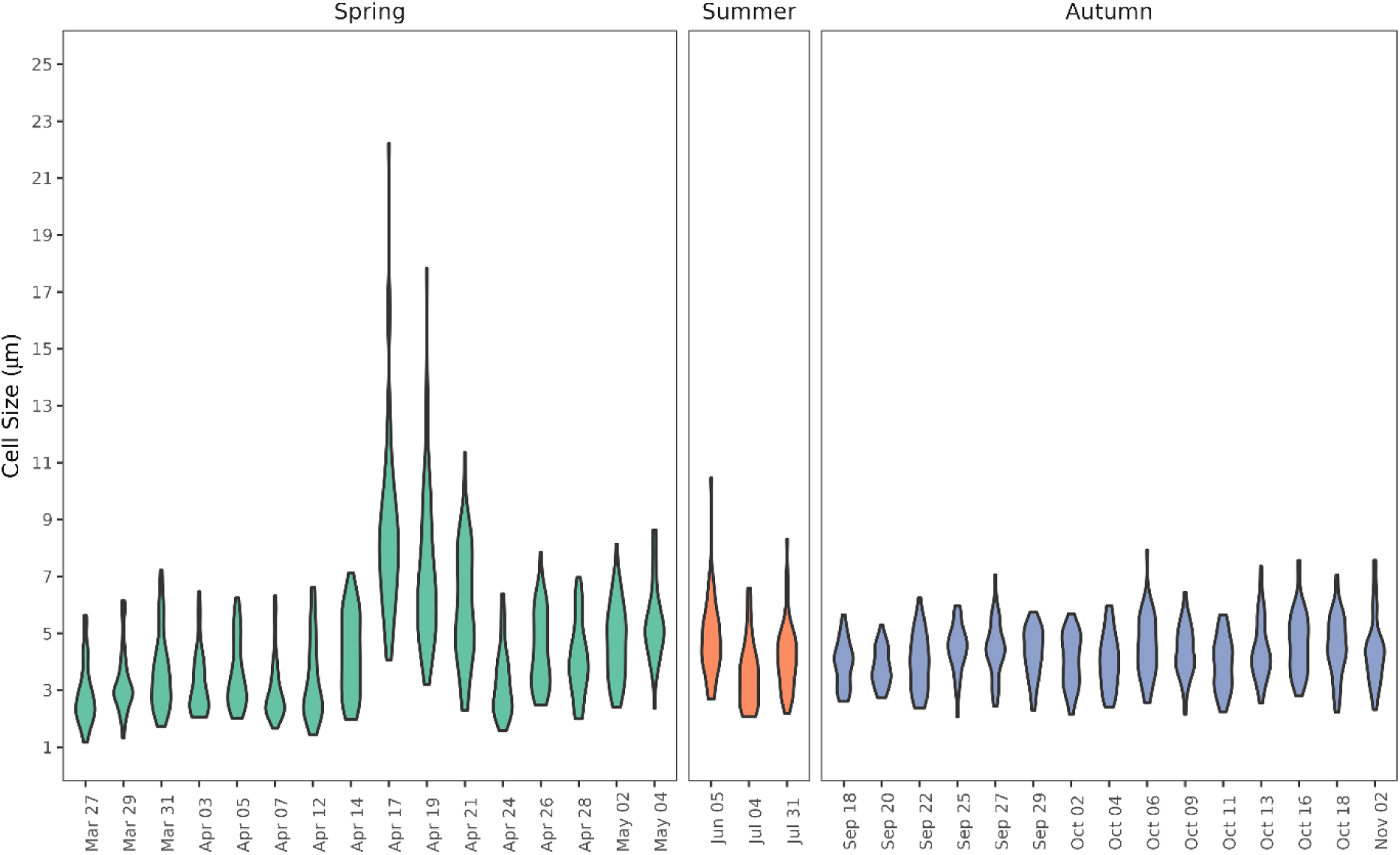
Violin plot of cell-size distribution of heterotrophic nanoflagellates (HNF) during high-frequency sampling campaigns in spring and autumn, and in monthly summer samplings. Fifty cells were measured per sample.

### Dynamics of HNF community composition

We used high-throughput sequencing of the V6-V8 region of the 18S rRNA gene to identify abundant groups of HNF. It showed that groups containing HNF contributed from <5 to > 20% of all eukaryotic reads, with Rhizaria (Cercozoa) being the most abundant (Supplementary Figure 2A). However, our previous attempt to quantify Cercozoa in the Baltic was unsuccessful (Piwosz et al., 2025). Therefore, we focused on other abundant groups such as MAST-1, MAST-2, and MAST-4 stramenopiles, *Paraphysomonas* spp., Katablepharidaceae, heterotrophic CRY1 cryptophytes and Telonemia for CARD-FISH analysis. PacBio long-amplicon sequencing was used for phylogenetic analysis to provide detailed insights to dynamics of the specific lineages within two most abundant HNF groups as revealed by CARD-FISH: katablepharids and MAST-2. Despite the total number of reads was 39% lower than from Illumina, the overall picture of the dynamics of HNF community was similar, with contribution of HNF groups to total read count ranging from 2 to 25% (Supplementary Figure 2B).

Bacterivorous HNF from groups MAST-1, MAST-4, CRY1 cryptophytes and *Paraphysomonas* ssp. contributed < 2% to the total HNF abundance. MAST-1 lineage was most abundant in June with the highest value recorded at 395 cells ml⁻¹, while being generally undetected or at very low levels from March to April (Figure 3A). Throughout the summer and autumn, the cell counts stayed low, ranging between 30 and 170 cells ml⁻¹. CRY1 was only detected in September, with the highest abundance recorded at 85 cells ml⁻¹, while being undetected during most of the seasonal period. *Paraphysomonas* ssp. abundance peaked at 80 cells ml⁻¹ in April and 260 cells ml⁻¹ in September, with low or undetectable levels in spring and autumn (Figure 3B).

**Figure 3.**
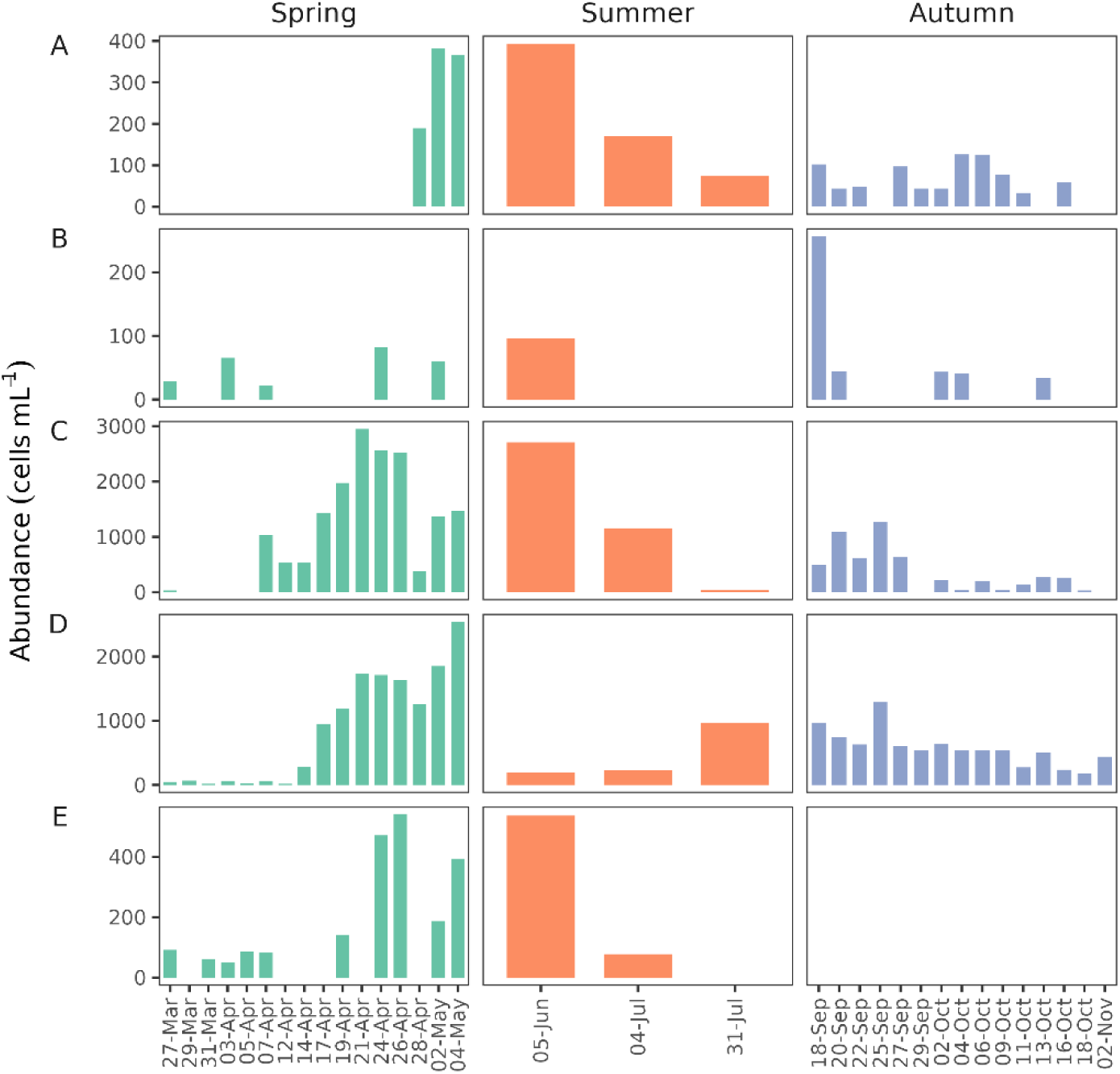
Abundances of HNF groups studied by CARD-FISH: MAST-1 stramenopiles (A); Paraphysomonas (B); katablepharids (C); MAST-2 stramenopiles (D); and Telonemia (E) during high-frequency sampling campaigns in spring and autumn, and in monthly summer samplings. Y-axis differs between the panels.

Omnivorous katablepharids were the most abundant HNF group, particularly in late April to early June. Their numbers increased sharply at the beginning of the sampling period, rising from negligible levels to 1,050 cells ml⁻¹ in the first week, then dropped by nearly 500 cells ml⁻¹ within the next two days then surged again to 1,430 cells ml⁻¹ and gradually raised to their peak of 2,945 cells ml⁻¹ in late April (Figure 3C). During the summer, the cell numbers varied reaching about 2,710 cells ml⁻¹ in early June but then dropped to approximately 1,160 cells ml⁻¹ in July. In autumn, the cell numbers ranged from 490 to 1,270 cells ml⁻¹ in September. By October, their abundances further decreased to 40 cells ml⁻¹, and katablepharids were no longer detected in November. Phylogenetic analysis of long amplicon sequencing provided detailed insights to the composition and dynamics of the specific katablepharids lineages (Figure 4). A group of highly similar ASV clustered as a sister branch to *Leucocryptos marina* with support > 93, likely representing separate species of this genus (referred to here as *Leucocryptos* sp.). Another set of *Leucocryptos*-related ASVs from a well-supported sister branch to *Leucocryptos marina* and newly defined *Leucocryptos* sp. We also identified three strongly supported clades within *Katablepharis* genus*: Katablepharis japonica, Katablepharis* sp.1 and *Katablepharis* sp.2 (Supplementary Figure 3). The latter comprise two ASVs weakly clustered (support <70) with *K. remigera.* These newly defined phylogroups showed remarkable dynamics. *Katablepharis* sp.1 and *Leucocryptos*-related ASVs dominated katablepharids community in spring, especially during the peak abundance, while *Katablepharis* sp.1 and *Leucocryptos* sp in autumn. *Katablepharis* sp.2 and *K. japonica* were detected in autumn, but they were less abundant (Figure 4A).

**Figure 4.**
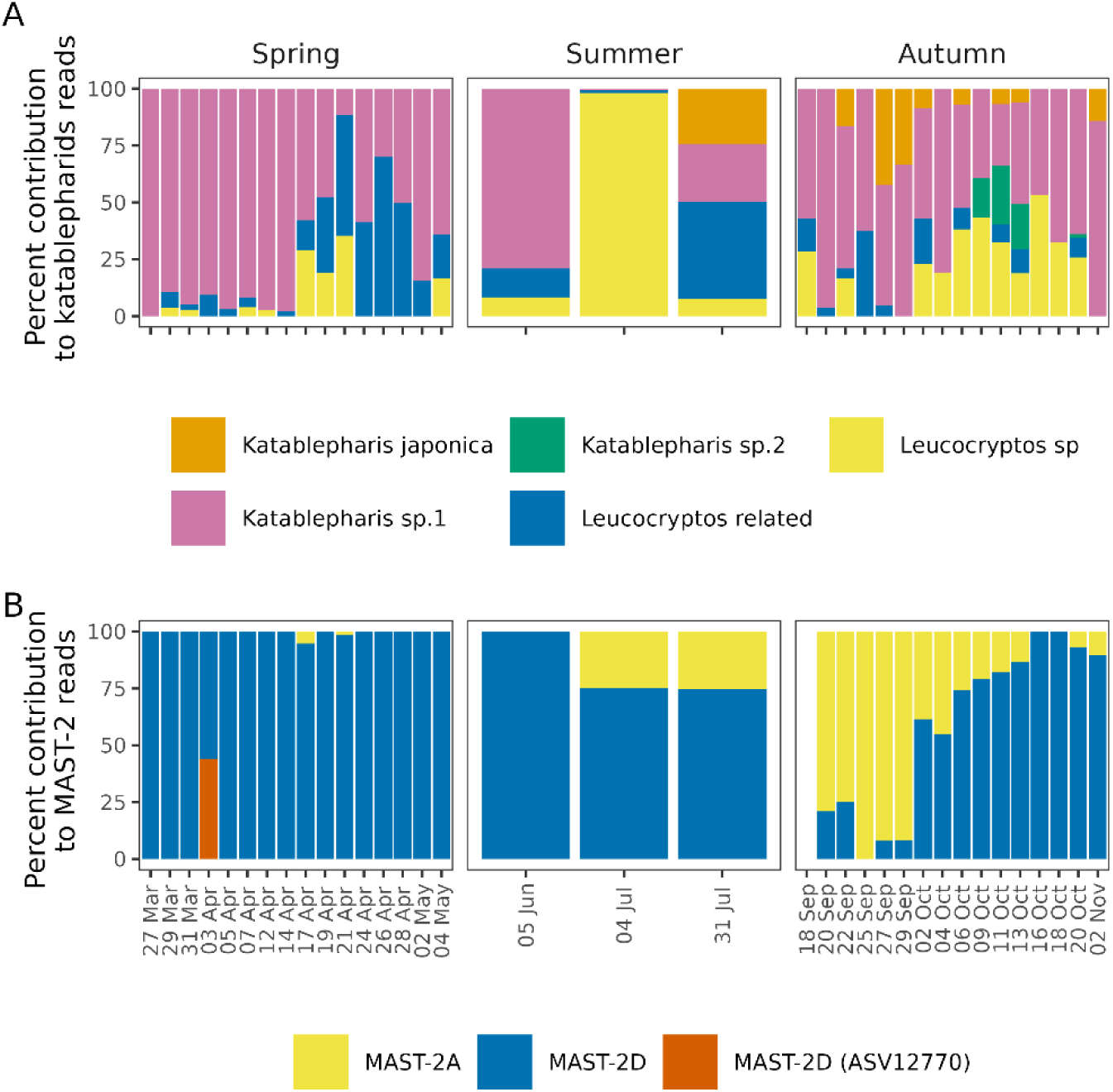
Composition of katablepharids (A) and MAST-2 (B) communities during high-frequency sampling campaigns in spring and autumn, and in monthly summer samplings from PacBio long amplicon sequencing. Bars show percent contribution of phylotypes identified based on phylogenetic analysis (Supplementary Figures 3 and 4).

MAST-2 peaked in late April and early May, with cell counts reaching approximately 2,500 cells ml⁻¹ (Figure 3D). After this peak, the numbers dropped sharply in June to about 190 cells ml⁻¹. Throughout the summer, the cell counts stayed lower, ranging between 200 and 1,000 cells ml⁻¹. Another smaller peak occurred in late September, with cell counts rising to over 1,200 cells ml⁻¹. By November, the numbers decreased again to about 430 cells ml⁻¹. Long amplicons sequencing revealed two main lineages of MAST-2 in our samples: MAST-2A and MAST-2D (Supplementary Figure 4). ASVs affiliated with MAST-2A were similar to each other and also to previously described sequences of MAST-2A lineage, forming a well-supported branch. In contrast, almost all MAST-2D-related ASVs formed a separate, well-supported branch within MAST-2D lineage, together with two other sequences. Only ASV12770 separated at the base of this branch (Supplementary Figure 4). MAST-2D contributed most to the read numbers during the spring peak, with ASV12770 detected only once during early spring (Figure 4B). MAST-2D contributed the most to the MAST-2 reads also in summer, although starting from July, contribution of MAST-2A increased, and this lineage dominated until end of September. From the beginning of October MAST-2D became more abundant and dominated MAST-2 community again (Figure 4B).

The abundance of Telonemia, a predatory group, varied from 50 to 540 cells ml⁻¹ between March and July, while it was mostly absent during the autumn months (Figure 3E).

### Microzooplankton

Ciliate abundance (excluding prevalently autotrophic *Mesodinium rubrum* and *M. major*) ranged from 3 to 71 cells ml ^-1^ throughout the sampling period. In spring, the highest abundance of 52 cells ml ^-1^ was recorded in April. In summer, abundance ranged from 12 to 19 cells ml ^-1^. In autumn, it varied from 4 to 71 cells ml^-1^, with the highest abundance recorded in September (Figure 5A). Their community was dominated by predatory and algivorous genera, such as flagellate predators *Urotricha* and *Balanion,* especially in spring and summer (Supplementary Figure 5). In autumn, there were two maxima of ciliate abundance (Figure 5A). The first one in late September was dominated by predatory *Balanion* and omnivorous *Lohmanniella*. The second peak in mid-October was dominated by omnivorous *Tintinnopsis*, *Tintinnidium, Lohmanniella*, *Rimostrombidium* and *Strombidium* in addition to *Balanion* (Supplementary Figure 5).

**Figure 5.**
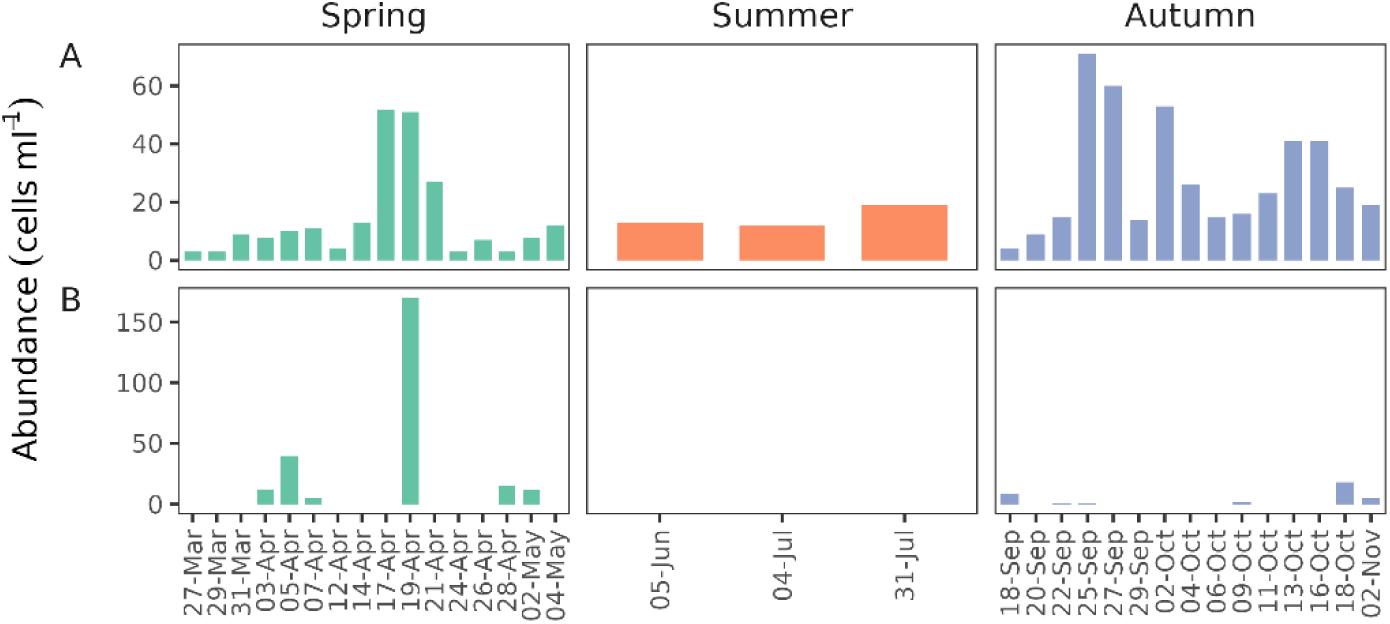
Abundance of heterotrophic ciliates (A) and heterotrophic dinoflagellates (B) during high-frequency sampling campaigns in spring and autumn, and in monthly summer samplings. Y-axis differs between the panels.

Heterotrophic dinoflagellates were present at only few occasions (Figure 5B). Their highest abundance of 170 cells ml ^-1^ was formed by algivorous Gymnodiniales and *Oblea rotunda* on April 19, during the peak of phytoplankton bloom. Except for this single maximum, their abundances were negligible, indicating heterotrophic dinoflagellates were not important predators in microbial food web during our study.

### Correlations Between Biological and Environmental Variables Across Seasons

Correlations were calculated for absolute abundance data of all HNF and of two most abundant groups, based on CARD-FISH: katablepharids and MAST-2. Temperature was a key environmental variable in our high frequency sampling campaigns both in spring and autumn (Figure 6, Supplementary Table 2). It displayed strong correlations with other environmental factors, with distinct seasonal patterns. In spring, temperature correlated positively with concentrations of DIN, N-NO_3_, N-NH_4_, and N:P ratio (Figure 6A). In contrast, autumn showed fewer significant correlations with temperature, with negative relationships being more prevalent (Figure 6, Supplementary Table 2). Notably, N:P ratio and N-tot exhibited strong negative correlations with temperature (Figure 6B). The environmental variables were also correlated with abundances of HNF and of katablepharids and MAST-2 revealed by CARD-FISH (Figure 6, Supplementary Table 2). In spring, temperature and DIN significantly correlated with all these flagellate groups; N-NH_4_ and P-PO_4_with HNF and MAST-2; N-NO_3_ with HNF and katablepharids; and N:P ratio with katablepharids (Figure 6A). In contrast, correlations were weaker in autumn, with temperature being the only environmental factor significantly positively correlated with HNF, katablepharids, and MAST-2, while N-tot and N:P Redfield ratio exhibited negative correlations, highlighting seasonal differences in environmental influences (Figure 6B). However, these relationships could be influenced by the strong correlation between temperature and other environmental variables. Therefore, to uncover potential dependencies masked by temperature, residual regression was applied for each season. While this approach allows to reveal additional correlations with other environmental variables, it actually eliminates factors strongly correlated with temperature. Consequently, there is a risk that significant environmental effects could be unintentionally excluded, limiting the ability to fully disentangle their influence through statistical methods. Surprisingly, these analyses indicated that P-PO_4_ was positively correlated with HNF in spring (r=0.63, p-value: 0.0094) and with MAST-2 in both seasons (spring: 0.63 r, p-value: 0.0089, autumn: r: 0.58, p-value:0.028, Supplementary Table 3). Analyses performed on the temporally shifted data at one- or two-interval lags also showed significant correlations, but there was no improvement in fit or interpretation compared to zero-lag data, indicating no delayed effects of environmental factors on biological organisms.

**Figure 6.**
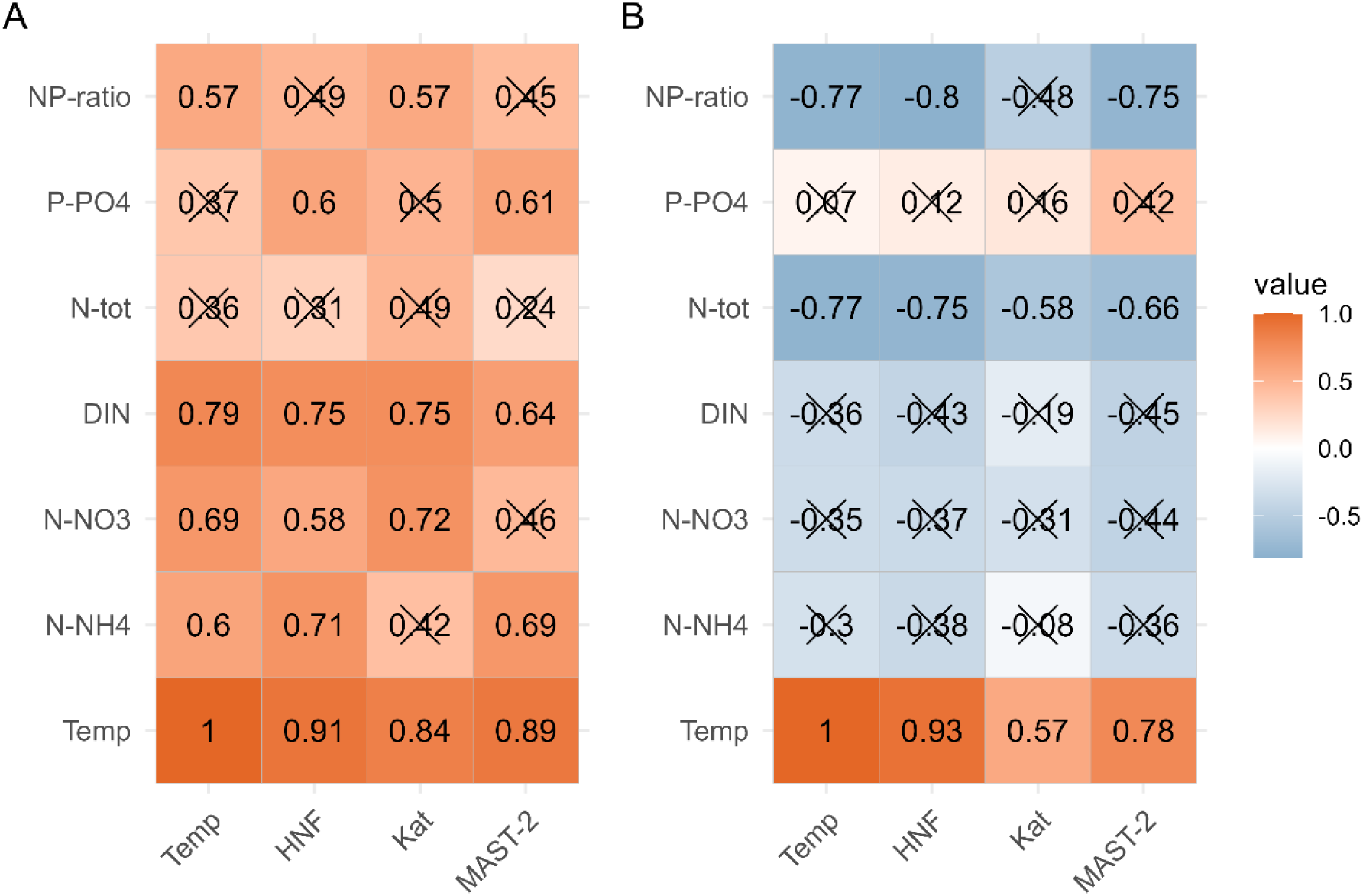
Correlation matrixes for the spring (A) and autumn high-frequency sampling campaigns showing values of Pearson correlation coefficients for relationships between environmental variables and the abundance of selected HNF groups studied by CARD-FISH. Non-significant correlations (p > 0.05) are crossed out. NP-ratio: ratio of concentration of total nitrogen (N-tot) to concentration of total phosphorus (P-tot); P-PO4: concentration of phosphate phosphorus; N-tot: concentration of total nitrogen; DIN: concentration of dissolved inorganic nitrogen; N-NO3; concentration of nitrate nitrogen; N-NH4: concentration of ammonium nitrogen; Temp: temperature; HNF: abundance of heterotrophic nanoflagellates; Kat: abundance of katablepharids, MAST-2: abundance of MAST-2 stramenopiles.

## Discussion

Microbial food webs play a critical role in the carbon flow and dynamics of aquatic ecosystems, especially during periods of phytoplankton blooms, which are common in temperate zone of the oceans, coastal areas, and freshwater environments (Trombetta et al., 2019), providing essential resources for aquatic food webs. Our high-frequency sampling, conducted during spring and autumn blooms in the Baltic Sea, provides insights into microbial food web dynamics with the focus on role of HNF. The key advancement of our study is the combination of CARD-FISH and long amplicon sequencing that allowed us to estimate the dynamics of narrower protist taxa by relating the absolute cell numbers detected by CARD-FISH probes to the relative contributions of more specific ASVs, such as specific subgroups shown in Fig. 4 (compare Figs. 3 and 4).

Eight eukaryotic groups in the Baltic Sea were quantified by CARD-FISH using specific probes to understand their response to environmental changes. The abundance of MAST-1 cells remained low throughout the study period, but a peak in June (Figure 3) suggests they may be adapted to summer conditions, even though they are associated with high-latitude regions (Obiol et al., 2024). On the other hand, their presence across different temperature ranges, including in the Mediterranean coastal environment (Massana et al., 2006a), suggests that while temperature influences their distribution, other environmental factors, such as nutrient availability and trophic interactions, likely contribute to their seasonal dynamics. CRY1 cryptophytes were below the detection limit in our samples, which differs from our previous CARD-FISH study in the Baltic, where they were consistently present and contributed considerably to the HNF community (Piwosz, 2019; Piwosz et al., 2025). This suggests that CRY1 cryptophytes exhibit substantial temporal and spatial variability in the Baltic Sea, potentially driven by competitive interactions, or grazing by predatory and omnivorous HNF and ciliates (Šimek et al., 2020). Variability in CRY1 abundance across ecosystems and time points has been reported in previous studies (Piwosz et al., 2016; Piwosz, 2019; Šimek et al., 2023), further highlighting their dynamic nature. *Paraphysomonas*, although widely distributed, consistently contributes little to HNF abundance, reflecting its specialized ecological role and an opportunistic life strategy, as emphasized in prior studies (Lim et al., 1999). In our study, it remained low or undetectable for most of spring and autumn, with brief peaks in April and September, indicating it flourishes briefly under favourable conditions. Katablepharids, as the second most abundant HNF group in our study, clearly play a dominant role in shaping microbial community dynamics in the Baltic Sea. Their sharp rise in abundance during the spring phytoplankton bloom suggests a strong dependence on nutrient-rich conditions and the availability of small or decaying algae and bacteria as prey (Kwon et al., 2017; Šimek et al., 2020). Their sustained abundance through early summer, followed by a decline in mid-summer and autumn, reflects the impact of rising temperatures, nutrient depletion, and seasonal shifts in microbial community composition, and likely also reflect shifts in prey availability. Long amplicon sequencing provided detailed insights into their diversity, revealing that while certain katablepharids types, such as *Katablepharis sp.1* and *Leucocryptos-related* species, remained dominant across seasons, other types, like *Katablepharis sp.2* and *Katablepharis japonica*, exhibited seasonal variations. This suggests that members of Katablepharidacea respond differently to environmental changes, which is also supported by the fact that correlations of katablepharids’ abundance with environmental variables varied seasonally (Figure 6).

The seasonal dynamics of MAST-2 further highlight the complex interactions within the microbial food web. MAST-2 is found in both freshwater and marine environments, especially in coastal and marine areas (Obiol et al., 2024). The peak abundance of MAST-2 during late spring and early summer coincided with the katablepharids (Figure 3), suggesting a potential predator-prey relationship between these groups. On the other hand, predator response to prey abundance is often delayed (Shabarova et al., 2021), thus we cannot exclude that the observed cooccurrence was at least partially driven by other factors, such as increase in picoplanktonic prey or response to environmental factors. The dominance of MAST-2D during spring and the initial dominance of MAST-2A in autumn, followed by MAST-2D (Figure 4) may explain differences in correlations of all MAST-2 with nutrients that were positive in spring and negative in autumn (Figure 6). However, we cannot exclude that these correlations with nutrients was driven by other, unstudied factors that differ between spring and autumn, most notably food preferences and community composition of prey. For instance, the smaller autumn peak in MAST-2 abundance suggests that their role extends into later stages of the microbial food web, potentially preying on other HNF groups or smaller protists as katablepharids declined. Telonemia, another predatory group, exhibited a distinct temporal pattern, with high abundance in spring and early summer and near absence by autumn. The absence of Telonemia in autumn contrasts with previous findings (Boukheloua et al., 2024), which may reflect differences in temperature preferences between the freshwater and brackish/marine-lineages, nutrient availability, or regional environmental conditions.

Size distribution of the HNF community affects the food web structure. Generally, there is little size variation within a single nanoflagellate species, while groups of multiple species show a broader size range (Piwosz, 2019). For instance, bacterivorous HNFs are typically very small (< 3 µm), while larger HNFs, such katablepharids or flagellated cercozoans are often omnivorous or predatory (Piwosz and Pernthaler, 2011; Grujcic et al., 2018; Šimek et al., 2020; Piwosz et al., 2021). The size distribution of HNF (Figure 2) agreed with the CARD-FISH results showing higher abundance of omnivorous katablepharids and predatory MAST-2 compared to bacterivorous groups (Figure 3). Concordantly, the cell size of majority of HNF was between 3 and 5 µm in all the seasons, suggesting they were omnivores rather than bacterivores (typically < 3 µm; Jürgens and Massana, 2008). The relatively low abundance of predatory HNF (>10) in autumn suggests a shift in the microbial food web, possibly with omnivorous HNF and ciliates playing a larger role in bacterial grazing (*e.g.* genus *Strombidium* and tintinnids).

We propose a refined model of the microbial food web in spring, defining the key HNF groups and their ecological roles based on our findings. The significant increase in total HNF abundance during this season, likely driven by favourable conditions such as temperature, bacterial growth boosted by phytoplankton biomass accumulation, and extracellular organic carbon release during bloom events (Zeder et al., 2009; Eckert et al., 2012), influences bacterial community structure, nutrient cycling, and energy transfer to higher consumers (Jezbera et al., 2006). These conditions lead to active trophic interactions, with bacterivorous and omnivorous HNF act as intermediates between bacteria and higher trophic levels, while predatory HNFs contribute to top-down regulation, reinforcing their role in microbial trophic interactions. However, we do not propose a refined microbial food web model for autumn, as reduced HNF abundance, lower phytoplankton biomass, and changing environmental conditions suggest different trophic interactions that are less understood and require further investigation.

The high variability of phytoplankton across both time and space complicates species-specific trend analyses, especially when sampling frequency is low (Wasmund and Uhlig, 2003). During spring (March to May), increasing temperatures and higher nutrient availability were generally linked to stronger phytoplankton blooms, with biomass reaching its highest levels in April. Ciliates, which peaked in abundance during the same period, are often considered potential grazers of phytoplankton (Sommer et al., 2012; Posch et al., 2015), although their exact role in regulating bloom dynamics, compared to large zooplankton, remains uncertain. In contrast, autumn (September to November) was characterized by a gradual cooling trend from warmer summer temperatures, accompanied by declining nutrient levels, which likely contributed to reduced phytoplankton growth. The gradual decline in HNF abundance during this period suggests that the shift in microbial interactions may be driven by competition for limited resources (such as nutrients) among microbial populations, rather than a sudden trophic shift. Microzooplankton, including typical flagellate hunters, such as ciliates from genera *Urotricha and Balanion* (Posch et al., 2015; Šimek et al., 2020), and dinoflagellates, likely influenced HNF abundance through direct grazing interactions along with a decreasing availability of bacterial prey. While *Urotricha* spp. and *Balanion* sp. are known predators of small algae and flagellates (Müller and Schlegel, 1999; Posch et al., 2015), their potential impact on HNF populations, documented also experimentally in freshwaters (Šimek et al., 2020, Asghar et al., in press), remains to be explored in the brackish environment. Overall, spring had stronger blooms due to favourable temperatures and nutrient availability, while autumn showed lower biomass with limited availability of nutrients along with the role of omnivorous ciliates triggering the decline of phytoplankton. Further high-temporal resolution studies are essential to enhance our understanding of the rapidly growing microbial populations during phytoplankton blooms, as there remains an ongoing debate regarding the influence of bottom-up and top-down factors on these events.

## Conclusions

This study emphasizes a key role of omnivorous and predatory HNF in the Baltic Sea microbial food web. Groups such as Katablepharids, MAST-2 and Telonemia were more abundant than typical bacterivores from CRY1 cryptophytes or MAST-1 and MAST-4 lineages, indicating that they may be major source of bacterial mortality in the coastal Baltic Sea. They are a link to mesozooplankton via predatory and omnivorous ciliates rather than heterotrophic dinoflagellates. High frequency sampling documented immense dynamics of all components of microbial food webs from primary producers (phytoplankton) via prokaryotes, HNF to microzooplankton. The changes in total abundances were accompanied by changes in community composition, as shown by combination of light microscopy, CARD-FISH, and long-amplicon sequencing. Further experimental work and more high frequency campaign are needed to fully understand dynamics of trophic interactions within microbial food webs.

## Acknowledgements

We thank Bartosz Witalis for helping with sample collection and Uroosa Uroosa for her assistance in sample filtration for DNA extraction. This project was funded by the National Science Centre, Poland under the Weave-UNISONO call in the Weave programme, project no. 2021/03/Y/NZ8/00076. I.M., T.S. and K. Š. were supported by the research grant 22-35826 K (Grant Agency of the Czech Republic) awarded to I.M.

## Author contributions

S.H.: Data curation, Formal analysis, Investigation, Visualisation, Writing – original draft; KW: Investigation; JC: Formal analysis, Visualisation, Writing – review and editing; KR: Investigation, Data curation, Writing – review and editing; LN: Investigation, Data curation; AA: Investigation, Data curation, Writing – review and editing; TS: Formal analysis, Writing – review and editing; IM: Conceptualization, Funding acquisition, Investigation, Writing – review and editing; KŠ: Conceptualization, Writing – review and editing; AJ: Investigation; MZ: Investigation: KP: Conceptualization, Data curation, Investigation, Funding acquisition, Project administration, Supervision, Visualisation, Writing – review and editing.

## Supplementary Information

**Supplementary Table 1**

List of oligonucleotide probes used for protistan lineages and hybridization conditions.

**Supplementary Table 2**

Values of Pearson coefficients and p-values for correlations between selected environmental and biological variables for spring and autumn high-frequency sampling campaigns.

**Supplementary Table 3**

Values of Pearson coefficients and p-values for residual regression (that excludes temperature effect) between selected environmental and biological variables for spring and autumn high-frequency sampling campaigns.

**Supplementary Figure 1.**

Values of key environmental parameters measured during high-frequency sampling campaigns in spring and autumn, and in monthly summer samplings. (A) Temperature (°C), (B) Salinity, (C) Concentration of Dissolved Inorganic Nitrogen (µmol dm⁻³), (D) Concentration of Total Nitrogen (µmol dm⁻³), (E) Concentration of Phosphate (µmol dm⁻³), and (F) Concentration of Total Phosphorus (µmol dm⁻³). Bars represent individual sampling dates, with colours indicating the corresponding season (Spring, Summer, or Autumn). A xlsx file containing these data is available at https://zenodo.org/ (DOI: 10.5281/zenodo.15411382).

**Supplementary Figure 2.**

HNF community composition during high-frequency sampling campaigns in spring and autumn, and in monthly summer samplings based on (A) short-amplicons from Illumina sequencing and (B) longamplicons from PacBio sequencing. Percentage contribution of ASV affiliated to groups known to contain HNF species. Groups further studied by CARD-FISH are shown in colours, while those unstudied by CARD-FISH in greyscale.

**Supplementary Figure 3.**

Consensus maximum likelihood tree of katablepharids based on the alignment of ASV sequences from PacBio sequencing. Bootstrap values are indicated at each node. Sequence labels represent both environmental ASVs and reference sequences. The scale bar indicates 0.08 substitutions per site. The tree in Newick format and the alignment are available at https://zenodo.org/ (DOI: 10.5281/zenodo.15411382).

**Supplementary Figure 4.**

Consensus maximum likelihood tree of MAST-2 stramenopiles based on the alignment of ASV sequences from PacBio sequencing. Bootstrap values are indicated at each node. Sequence labels represent both environmental ASVs and reference sequences. The scale bar indicates 0.05 substitutions per site. The tree in Newick format and the alignment are available at https://zenodo.org/ (DOI: 10.5281/zenodo.15411382).

**Supplementary Figure 5.**

Ciliate community composition during high-frequency sampling campaigns in spring and autumn, and in monthly summer samplings based on light microscopy counts. A xlsx file containing these data and trophic roles of ciliate species is available at https://zenodo.org/ (DOI: 10.5281/zenodo.15411382).

