## Supplementary Figure 1 for "High-temporal resolution of microbial food web dynamics and structure during phytoplankton blooms in the Baltic Sea"

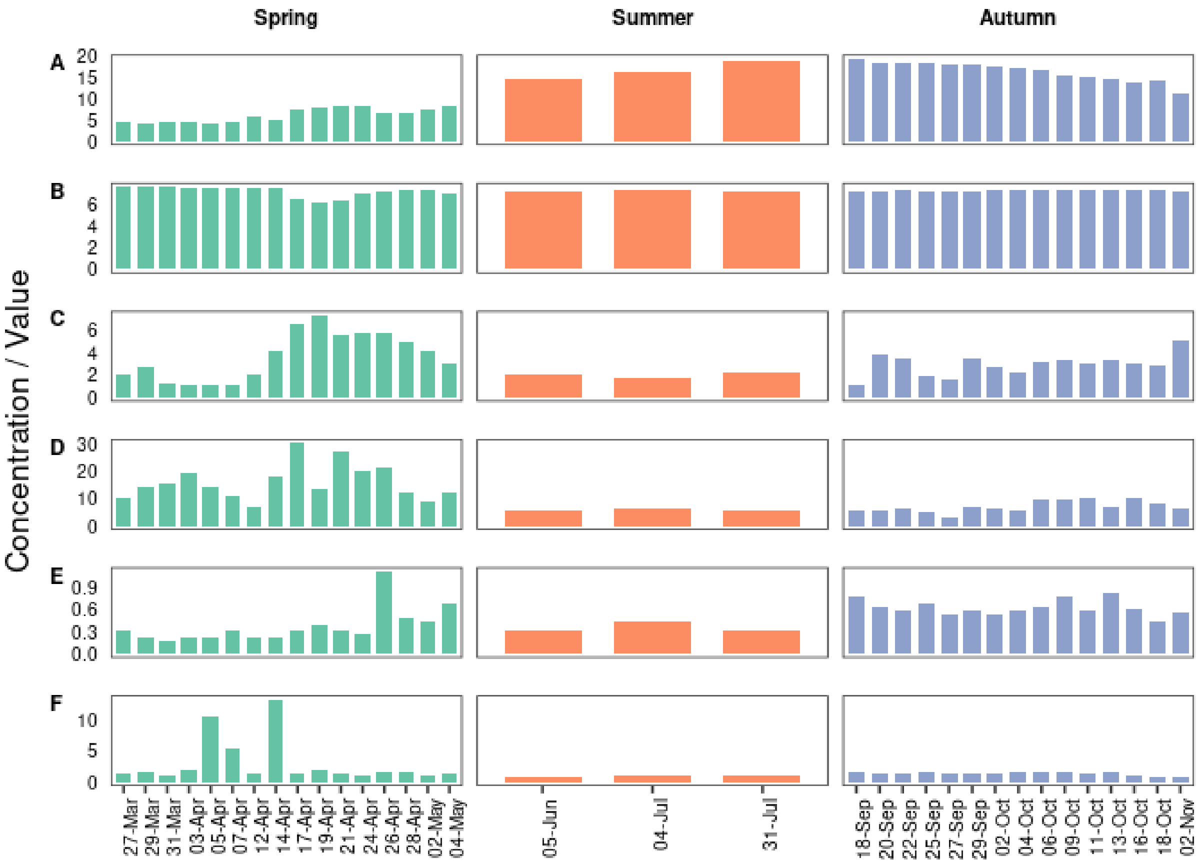


**Supplementary Figure S1**

Values of key environmental parameters measured during high-frequency sampling campaigns in spring and autumn, and in monthly summer samplings. (A) Temperature (°C), (B) Salinity, (C) Concentration of Dissolved Inorganic Nitrogen (µmol $dm^{-3}$), (D) Concentration of Total Nitrogen (µmol $dm^{-3}$), (E) Concentration of Phosphate (µmol $dm^{-3}$), and (F) Concentration of Total Phosphorus (µmol $dm^{-3}$). Bars represent individual sampling dates, with colours indicating the corresponding season (Spring, Summer, or Autumn). A xlsx file containing these data is available at https://zenodo.org/ (DOI: 10.5281/zenodo.15411382).
