## Supplementary Figure 2 for "High-temporal resolution of microbial food web dynamics and structure during phytoplankton blooms in the Baltic Sea"

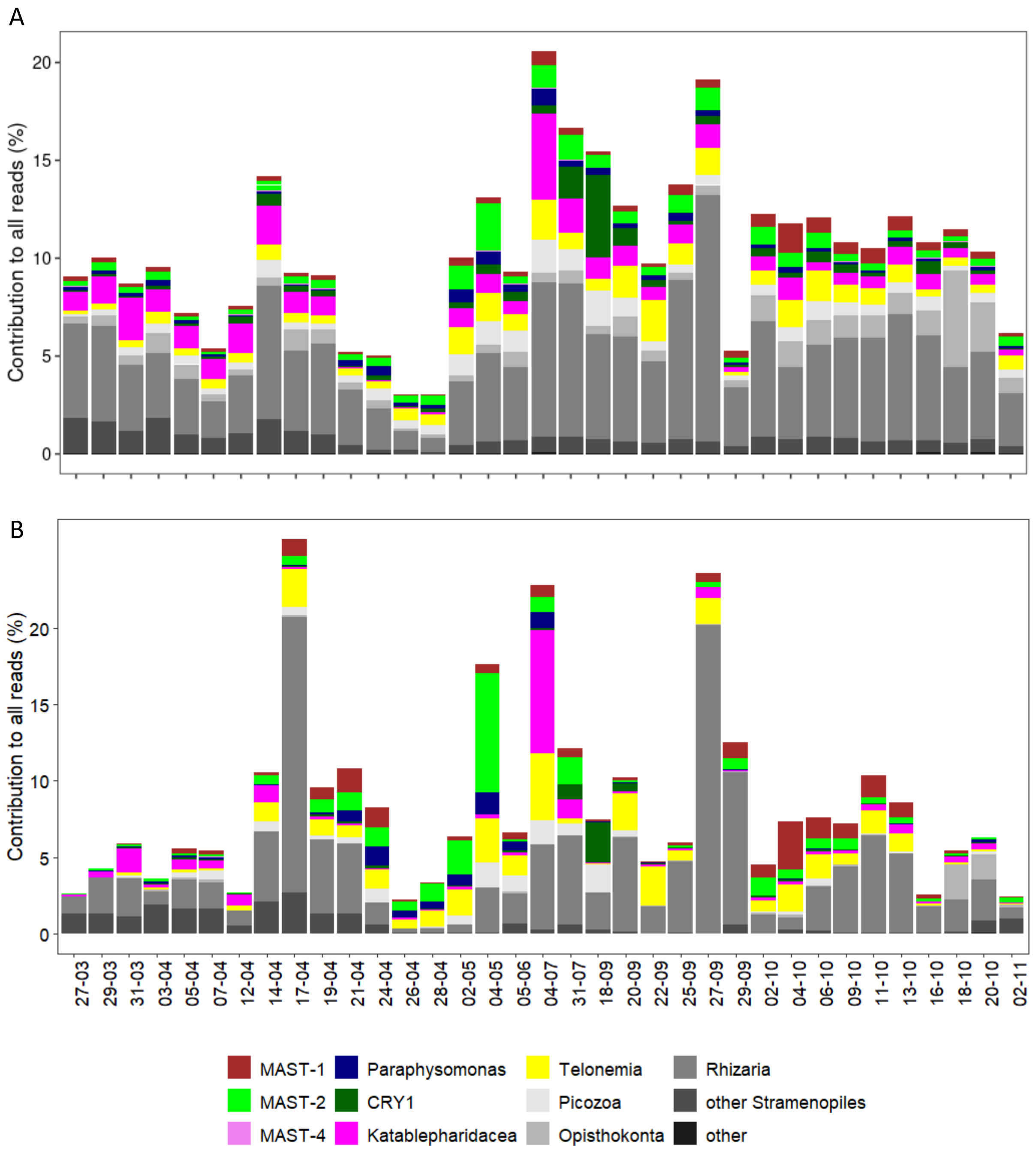


**Supplementary Figure S2**

HNF community composition during high-frequency sampling campaigns in spring and autumn, and in monthly summer samplings based on (A) short-amplicons from Illumina sequencing and (B) long-amplicons from PacBio sequencing. Percentage contribution of ASV affiliated to groups known to contain HNF species. Groups further studied by CARD-FISH are shown in colours, while those unstudied by CARD-FISH in greyscale.
