## Supplementary Figure 3 for "High-temporal resolution of microbial food web dynamics and structure during phytoplankton blooms in the Baltic Sea"

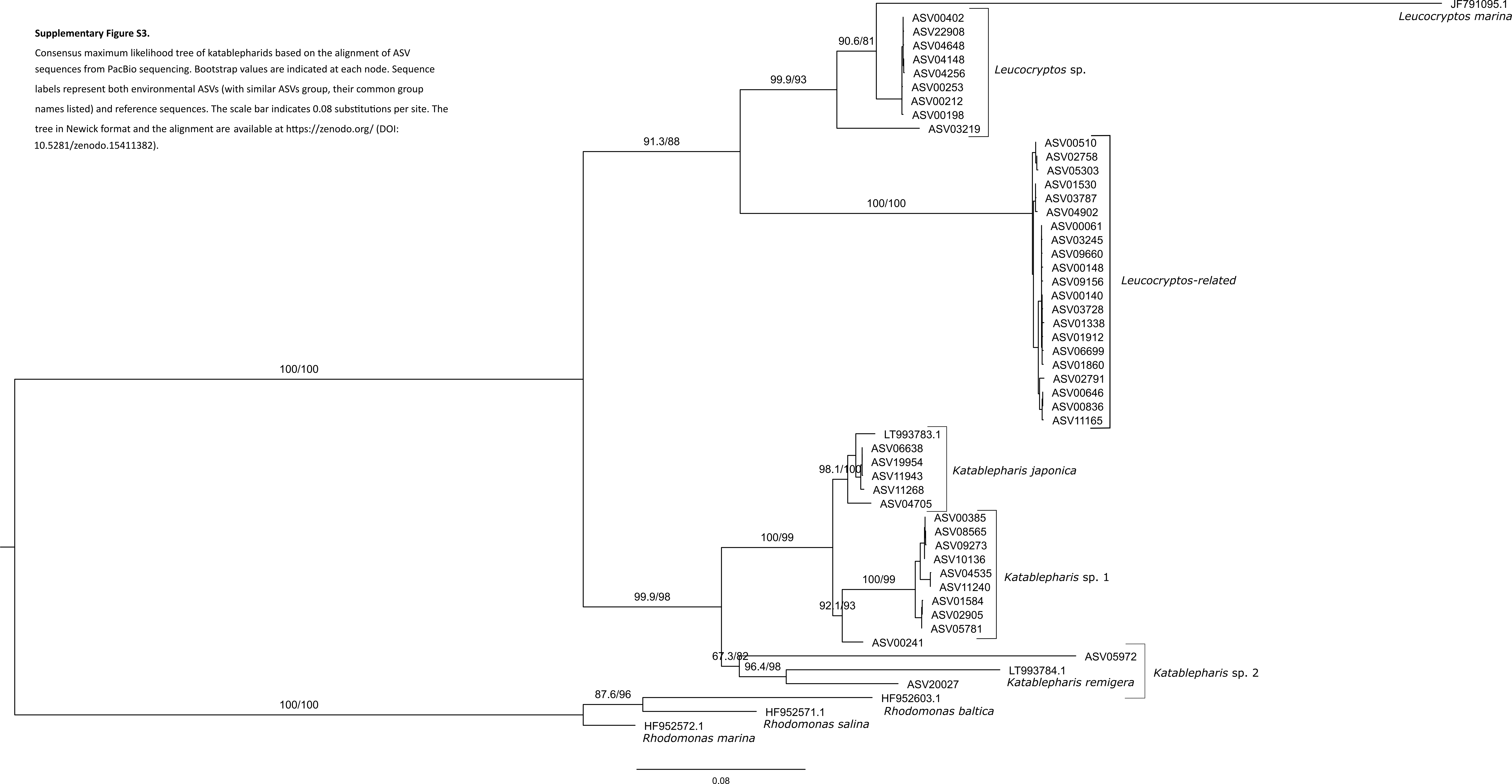


**Supplementary Figure S3.**

Consensus maximum likelihood tree of katablepharids based on the alignment of ASV sequences from PacBio sequencing. Bootstrap values are indicated at each node. Sequence labels represent both environmental ASVs (with similar ASVs group, their common group names listed) and reference sequences. The scale bar indicates 0.08 substitutions per site. The tree in Newick format and the alignment are available at https://zenodo.org/ (DOI: 10.5281/zenodo.15411382).
