## Supplementary Figure 5 for "High-temporal resolution of microbial food web dynamics and structure during phytoplankton blooms in the Baltic Sea"

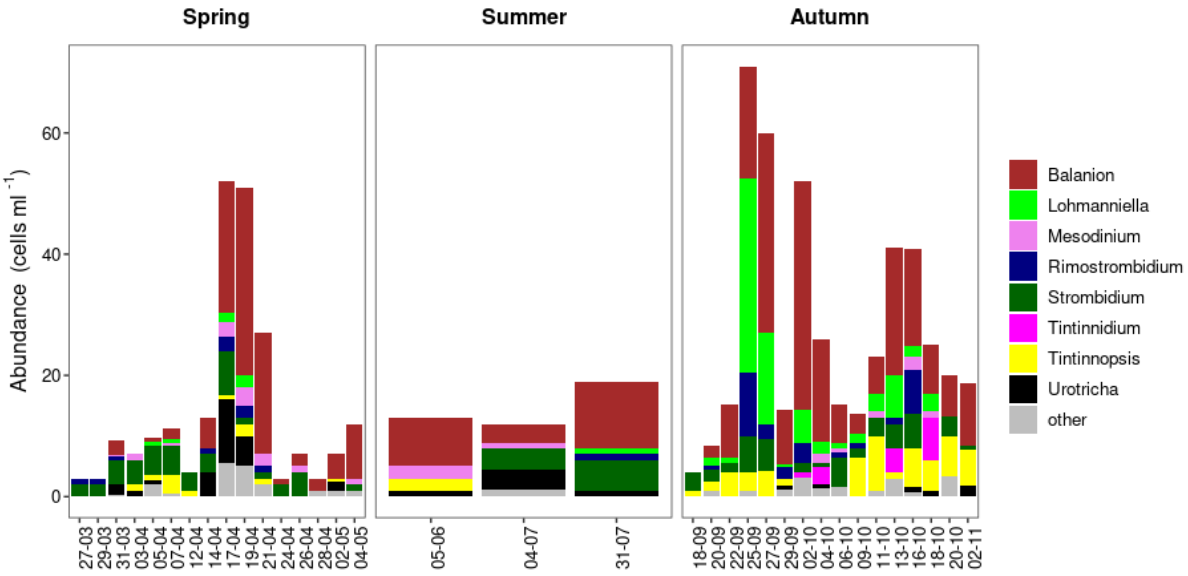


**Supplementary Figure S5.**

Ciliate community composition during high-frequency sampling campaigns in spring and autumn, and in monthly summer samplings based on light microscopy counts. A xlsx file containing these data and trophic roles of ciliate species is available at https://zenodo.org/ (DOI: 10.5281/zenodo.15411382).
