## Supplementary Table 1 for "High-temporal resolution of microbial food web dynamics and structure during phytoplankton blooms in the Baltic Sea"

**Supplementary Table 1.** List of oligonucleotide probes used for protistan lineages and hybridization conditions. % HB – the percentage of formamide in the hybridization buffer at the specified temperature (Temp).

| Probe name | Target group | References | Sequence (5' – 3') | % HB | Temp (°C) |
|---|---|---|---|---|---|
| **EUK-516** | Eukaryotes | (Amann *et al.*, 1990) | ACCAGACTTGCCCTCC | 20 | 35 |
| **NS2** | Marine stramenopiles lineage 2 | (Massana *et al.*, 2006) | ATGGGCCGACCGGTCGCT | 30 | 46 |
| **NS4** | Marine stramenopiles lineage 4 | (Massana *et al.*, 2002) | TACTTCGGTCTGCAAACC | 30 | 46 |
| **KAT900*** | Katablepharidacea | (Piwosz *et al.*, 2025) | ATAAACGCCCCCAACTATCC | 55 | 35 |
| **KAT900_C1** | | | ACAAACGCCCCCAACTATCC | | |
| **KAT900_C2** | | | ATTAACGCCCCCAACTATCC | | |
| **KAT900_C3** | | | ATGAACGCCCCCAACTATCC | | |
| **KAT900_C4** | | | ATAAATGCCCCCAACTATCC | | |
| **NS1** | Marine stramenopiles | (Massana *et al.*, 2004) | ATTACCTCGATCCGCAAA | 30 | 46 |
| **PBIV 1537** | *Paraphysomonas* | (Caron *et al.*, 1999) | CAAGATTCAGAATTGCAAAA A | 20 | 35 |
| **TELO-1250** | Telonemia | (Mukherjee *et al.*, 2024) | CAGYCAAGGTGGACAACTY GTT | 40 | 35 |
| **CRY1** | Cry1 lineage of cryptophytes | (Piwosz *et al.*, 2016) | CACAGTAAACGATCCGCGCA A | 40 | 35 |

* Probe Kat900 must be used with four competitors (Kat900C1-C4) to ensure specificity.
